# Enriched-GF: A Reproducible High-Yield Autologous Blood-Derived Growth Factor Method for Regenerative Medicine

**DOI:** 10.64898/2026.03.19.712883

**Authors:** Himanshu Bansal, Mahima Singhal, Alnkrita Bansal, Anupama Bansal, Shahnawaz Hussein Khan, Jerry Leon, Mustafa al Maini, Matias Fernandez Viña, Leonid Reyfman

## Abstract

**Background:** Platelet-derived Growth factors play key roles in tissue repair and regeneration, yet conventional platelet-rich plasma (PRP) formulations release these mediators inconsistently in vivo due to variability in platelet yield and activation dynamics. To overcome this limitation, direct administration of concentrated platelet-derived growth factor preparations has gained interest, though current manufacturing approaches for human platelet lysate (hPL), growth factor concentrates (GFC), and conditioned serum remain constrained by batch variability, incomplete platelet degranulation, and reliance on anticoagulants. Here, we examine alternative platelet activation workflows to establish a standardized, efficient, and reproducible method for high-yield growth factor recovery suitable for translational and clinical applications.

**Methods:** Nine GFC production protocols were compared, employing different combinations of freeze–thaw (FT) cycling, glass bead (GB) agitation, calcium (Ca^2^) activation, and a novel Enriched Growth Factor (Enriched-GF) method. The objective was to identify a protocol capable of maximizing growth factor yield within a three-hour workflow. Optimal Ca^2^ concentrations and GB conditions were determined from prior optimization studies and integrated into the Enriched-GF processing scheme. Platelet concentrates (n = 10 per protocol) were processed under each condition, and growth factor levels were quantified using ELISA.

**Results:** Growth factor yields differed significantly across protocols. The greatest and most consistent increases in growth factor release were observed with the Enriched-GF method combining GB activation, FT cycling, and Ca^2^ stimulation. This approach resulted in markedly elevated concentrations of key regenerative mediators, including enhanced EGF release, a 4.5-fold increase in PDGF, maximal TGF-β liberation, and a four-fold increase in FGF2 relative to conventional platelet lysate or conditioned serum preparations. These results were reproducible across independent donor pools, demonstrating robustness and batch-to-batch consistency.

**Conclusion:** We describe a rapid and reproducible method for producing highly concentrated platelet-derived growth factors using a combined GB–FT–Ca^2^ activation strategy. The Enriched-GF protocol consistently outperformed existing platelet lysate, conditioned serum, and conventional GFC preparation methods, yielding a standardized product with enhanced growth factor content. This Enriched-GF approach offers a clinically practicable solution for applications in regenerative medicine requiring reliable and high-yield growth factor delivery.

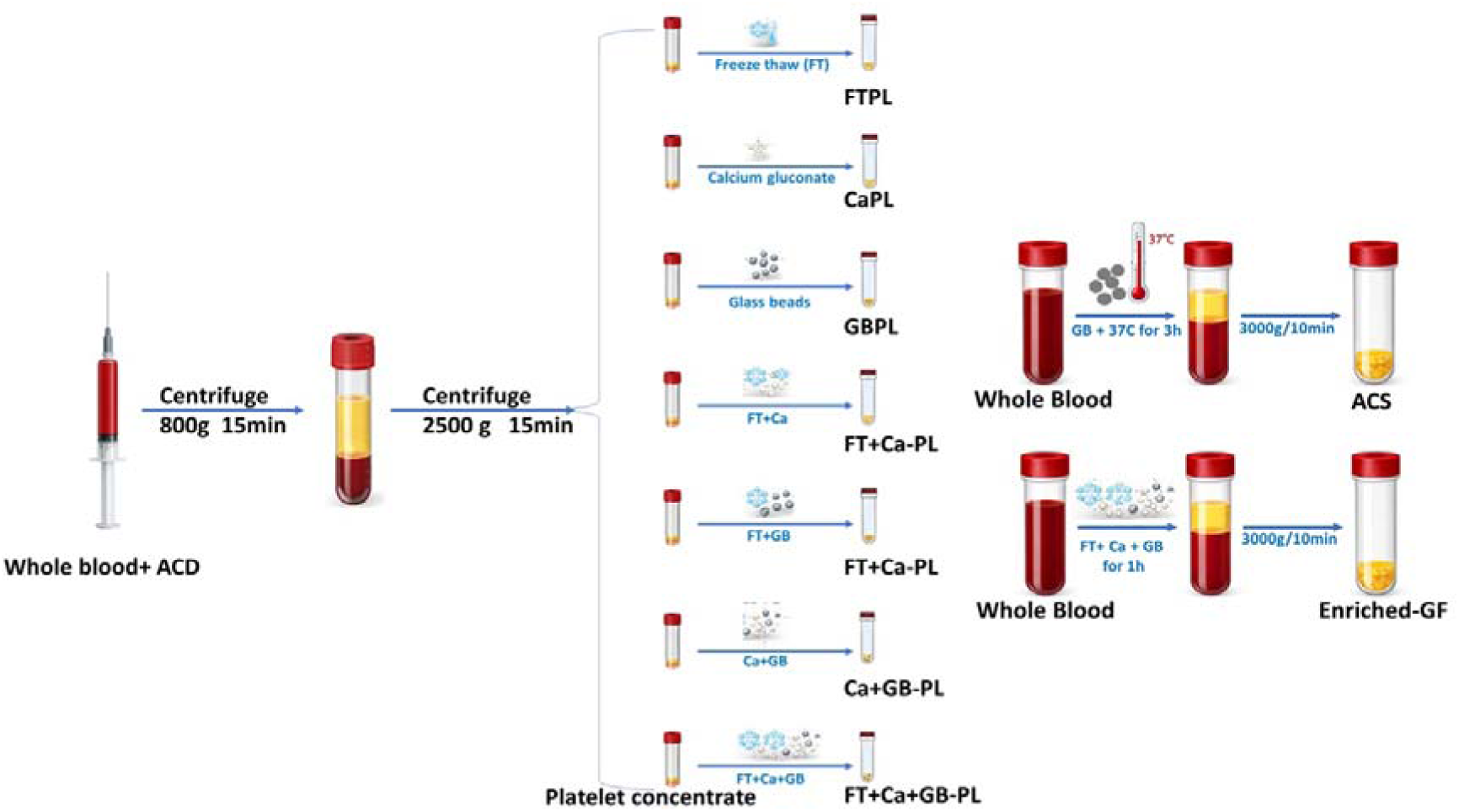

Schematic overview of platelet concentrate preparation from whole blood and the generation of different platelet lysates and growth factor-enriched serum using freeze-thaw, calcium gluconate, and glass bead activation methods.

## Introduction

Platelets are anucleate, discoid-shaped cells containing three primary types of secretory granules: lysosomes, dense granules, and α-granules ^1^. These granules enable platelets to execute a variety of biological roles beyond hemostasis, such as triggering and spreading inflammation ^1^, showing anti-inflammatory responses ^2–4^, and having the analgesic potential. Among them, the most common are alpha-granules. They are important stores of cytokines, growth factors, which promote tissue repair ^1, 5^, regulate inflammation and promote anabolic events, by recruiting and stimulating other inflammatory cells ^2, 3^. Antimicrobial effects have also been demonstrated by human platelet-derived products, though the mechanisms have not been completely understood ^6, 7^, Complement proteins and complement-binding proteins within α-granules may contribute to this effect ^8^.This and these few functions demonstrate that platelets are not merely important in the prevention of bleeding but also in the regulation of repair and immunological responses, and so platelets are good candidates to be used in regeneration.

Platelet-derived growth factor (PDGF) and vascular endothelial growth factor (VEGF) are some of the key growth factors that are contained in the alpha-granules and that are central to the process of wound healing and tissue regeneration ^1, 9, 10^. These factors promote tissue repair ^11^, attract mesenchymal cells ^11, 12^, stimulate extracellular matrix (ECM) production ^13^, and drive angiogenesis ^14, 15^. Accordingly, PRP has gained widespread clinical use over the past two decades. However, PRP performance remains inconsistent due to variability in platelet yield, activation state, and the incomplete or variable release of stored growth factors at the target site. These limitations have prompted growing interest in the direct administration of concentrated growth factor products, rather than relying on in vivo platelet activation. Recombinant growth factors have been used therapeutically; however, their high production costs and limited accessibility restrict broader clinical application ^16–18^. Using recombinant growth factors regularly is still not practicable. This makes the need for more accessible, biologically produced options even stronger.

As an alternative, hPL has emerged as an acellular derivative, a cost-effective, biologically active substitute that provides a rich source of growth factors, cytokines, chemokines, and osteoconductive proteins for cell therapy, tissue engineering, and regenerative medicine ^19–24^. Furthermore, HPL enables wider allogeneic use, facilitates batch pooling to reduce donor-to-donor variability, and enhances product standardisation ^21, 24^. It can be prepared from platelet concentrates using various methods such as manual centrifugation, plateletpheresis, or commercially available systems utilising centrifugation or filtration ^25^. Platelet activation to induce granule release can be achieved through freeze/thaw cycles ^26–28^, addition of thrombin ^29, 30^ or calcium chloride ^30^, or sonication ^31–33^, or solvent/detergent treatment ^25, 34^.The manner of activation, on the other hand, has a direct effect on the profile and concentration of the growth factors that are produced, which makes a big difference between preparations.

Despite increasing clinical interest, the heterogeneity of available methods results in significant variability in growth factor yields, limiting reproducibility and clinical translation. Commercial kits and proprietary protocols claim to maximise growth factor concentration, yet the actual levels of bioactive factors derived from each blood sample remain unpredictable. Consequently, hPL has delivered inconsistent therapeutic outcomes, particularly in pain management and tissue regeneration. Moreover, hPL is limited by its poor stability, with rapid loss of potency upon minimal storage. This lack of standardisation makes it harder to compare studies and makes it harder for regulators to approve and for doctors to use the results in their own work. Consequently, there exists an urgent need for a standardised, repeatable, and therapeutically dependable technique of hPL production that guarantees both stability and uniform bioactivity. However, many existing preparations contain intact cellular components or exhibit variability arising from platelet concentrate preparation and activation, which can result in incomplete growth factor release, injection-site discomfort, or potential immunogenic responses. To address these limitations, we investigated the Enriched-GF method, a next-generation, blood-derived growth factor formulation designed to isolate regenerative protein factors while remaining fully acellular. This approach enables enhanced and sustained growth factor release, preserves autologous origin, minimizes inflammatory responses, reduces the variability commonly associated with conventional preparations, and eliminates the need for xenogenic anticoagulants, thereby improving consistency and therapeutic outcomes. Collectively, Enriched-GF aims to retain the regenerative efficacy of conventional biologics while improving safety, reliability, and patient tolerability.

## Materials and Methods

### Study Approval and Donor Recruitment

This study was conducted in accordance with the ethical standards of the Anupam Hospital, Rudrapur. All procedures involving human participants were approved by the Institutional Ethical Committee. Written informed consent was obtained from all healthy adult donors before blood collection.

### Blood Collection

A total of 90 mL of venous blood was collected from each donor. Of this, 70 mL was drawn into sodium citrate anticoagulant tubes (blood: citrate = 9:1) for platelet concentrate (PC) preparation, while the remaining 20 mL was collected into tubes without anticoagulant for whole blood–derived lysate protocols. Samples were processed within 1 hour of collection.

### Generation of Platelet Concentrates (PCs)

PCs were prepared using a standard double-spin method. Briefly, 10 mL of citrate-anticoagulated whole blood was centrifuged at 800 rpm for 15 min at room temperature without brake (Eppendorf 5810R, Eppendorf AG, Hamburg, Germany). The plasma and buffy coat layers were transferred and centrifuged at 4,500 rpm for 15 min with the brake applied. The resulting platelet pellet was resuspended in 2 mL of autologous plasma. Platelet counts were verified to ensure a final concentration of ≥1 × 10□ platelets/mL before lysate preparation.

### Preparation of Platelet Lysates

PCs were processed using seven distinct activation strategies:

**1. Freeze–Thaw Platelet Lysate (FTPL)**

PCs were frozen at –80 °C and thawed at 37 °C repeated 3 cycles. Samples were centrifuged at 6,000 × g for 10 min, and the supernatant was filtered (0.22 µm) and aliquoted.

**2. Calcium-Activated Platelet Lysate (CaPL)**

PCs were mixed with calcium gluconate (1:10, v/v) and incubated for 1 h at room temperature. After clot retraction, samples were centrifuged at 6,000 × g for 10 min, filtered (0.22 µm), and aliquoted.

**3. Glass Bead–Activated Platelet Lysate (GB-PL)**

Sterile glass beads (0.5 g/mL) were added to PCs and incubated at 37 °C under gentle agitation until fibrin clot formation (∼60 min). Samples were centrifuged at 6,000 × g for 10 min, filtered (0.22 µm), and aliquoted.

**4. Calcium + Glass Bead Activation (Ca+GB-PL)**

PCs were mixed with calcium gluconate (1:10, v/v) and glass beads (0.5 g/mL) and incubated at room temperature for 1h. After clot formation, samples were centrifuged at 6,000 × g for 10 min, filtered (0.22 µm), and aliquoted.

**5. Freeze–Thaw + Glass Bead Activation (FT+GB-PL)**

PCs were combined with glass beads (0.5 g/mL), frozen at –80 °C for 1 h, thawed at 37 °C, centrifuged at 6,000 × g for 10 min, filtered (0.22 µm), and aliquoted.

**6. Freeze–Thaw + Calcium Activation (FT+CaPL)**

PCs were mixed with calcium gluconate (1:10, v/v), frozen at –80 °C for 3 h, thawed at 37 °C, centrifuged at 6,000 × g for 10 min, filtered (0.22 µm), and aliquoted.

**7. Freeze–Thaw + Calcium + Glass Bead Activation (FT+Ca+GB-PL)**

PCs were mixed with calcium gluconate (1:10, v/v) and glass beads (0.5 g/mL), frozen at –80 °C for 1 h, thawed at 37 °C, centrifuged at 6,000 × g for 10 min, filtered (0.22 µm), and aliquoted.

### Whole Blood–Derived GF Preparations Activated Conditioned Serum (ACS)

To generate ACS, whole blood (10 mL per donor) was collected into non-anticoagulated glass tubes pre-loaded with sterile borosilicate beads (approximately 0.5 g/mL blood). The tubes were incubated at 37 °C for 1 h to allow contact activation of coagulation and leukocyte-mediated cytokine release. Following incubation, the samples were centrifuged at 6,000 × g for 10 min. The resulting serum fraction was carefully collected, passed through a 0.22 µm sterile filter, aliquoted, and stored at –80 °C for subsequent growth factor quantification.

### Enriched-GF Method (WB-FT+Ca+GB)

Whole blood (10 mL) without anticoagulant was centrifuged at 1,000 × g for 15 min without a brake. Approximately 3 mL of plasma above the buffy coat was retained and then mixed with calcium gluconate (1:10, v/v) and glass beads (0.5 g/mL), frozen at –80 °C for 1 h, and thawed at 37 °C. After centrifugation at 6,000 × g for 10 min, the supernatant was filtered (0.22 µm), aliquoted. This preparation is designated Enriched-GF.

### Growth Factor and Cytokine Analysis of Platelet Lysate

The concentrations of soluble proteins in platelet lysate were analyzed using commercially available kits. Specifically, TGF-β1, PDGF, IGF-1, EGF, VEGF, bFGF, HGF were analyzed by enzyme-linked immunosorbent assay (ELISA) according to the manufacturer’s instructions (R&D Systems, Inc., Minneapolis, MN). ELISAs were performed in duplicates, and absorbance was read at 450 nm on a BioTek ELx808 ELISA plate reader (Agilent Technologies, USA).

## Results

Quantitative analyses of platelet-derived growth factors have revealed significant variations in growth factor release, contingent upon the activation strategy employed. Across all protocols, growth factor releases progressively increased with the complexity of activation, ranging from single-mode (FT, Ca^2^, or GB) to dual-mode (FT+CaPL, FT+GB-PL, or Ca+GBPL), and culminating in triple-mode activation (FT+Ca+GB-PL). Notably, the Enriched-GF protocol, which integrates sequential mechanical, thermal, and calcium-mediated activation, consistently yielded the highest concentrations of all measured growth factors with reduced inter-donor variability, indicative of more complete platelet degranulation and enhanced reproducibility. This enhanced activation likely facilitates more efficient platelet membrane disruption, promoting the release of a broader spectrum of growth factors.

Among the growth factors involved in epithelial regeneration and extracellular matrix remodeling, EGF exhibited a significant increase from 0.70 ng/mL in ACS to 1.69 ng/mL in Enriched-GF (P < 0.001) **(Fig. 1a)**. FGF-2 increased markedly from 0.15 ng/mL to 1.17 ng/mL, representing an approximate sevenfold enhancement (P < 0.001) **(Fig. 1b)**. IGF-1, recognized for its role in promoting anabolic tissue repair, also attained its signifcantly highest concentration in Enriched-GF (74.19 ng/mL) from 36.4ng/ml in ACS, exceeding levels observed in FT and Ca-only preparations (Fig. 1c). IGF-1, known for its role in facilitating anabolic tissue repair, achieved its significantly highest concentration in the Enriched-GF preparation (74.19 ng/mL), compared to 36.4 ng/mL in the autologous conditioned serum (ACS), surpassing the levels observed in FT and Ca-only preparations **(Fig. 1c)**.These elevations in EGF, FGF-2, and IGF-1 substantiate that regenerative signaling mediators are preferentially released when membrane disruption and coagulation pathways are sequentially activated, rather than individually. Growth factors predominantly stored within platelet α-granules demonstrated even more pronounced increases. PDGF, a principal chemoattractant and modulator of fibrosis, significantly rose from 67.025 ng/mL (FT+GB+Ca-PL) to 160.73 ng/mL in Enriched-GF, representing the largest fold-change observed (P < 0.001) **(Fig. 1d)**. Similarly, TGF-β increased substantially from 16.07 ng/mL in ACS to 80.42 ng/mL in Enriched-GF (P < 0.001), indicating highly efficient granule discharge under multi-modal activation **(Fig. 1e)**. VEGF, the principal angiogenic factor, also demonstrated a significant increase (P < 0.001) to 2.18 ng/mL in Enriched-GF compared with 0.69–1.62 ng/mL in single- or dual-mode activation methods **(Fig. 1f)**, suggesting that vascular signaling components are particularly responsive to the cumulative activation environment rather than to any singular stimulus.

**Figure 1.**
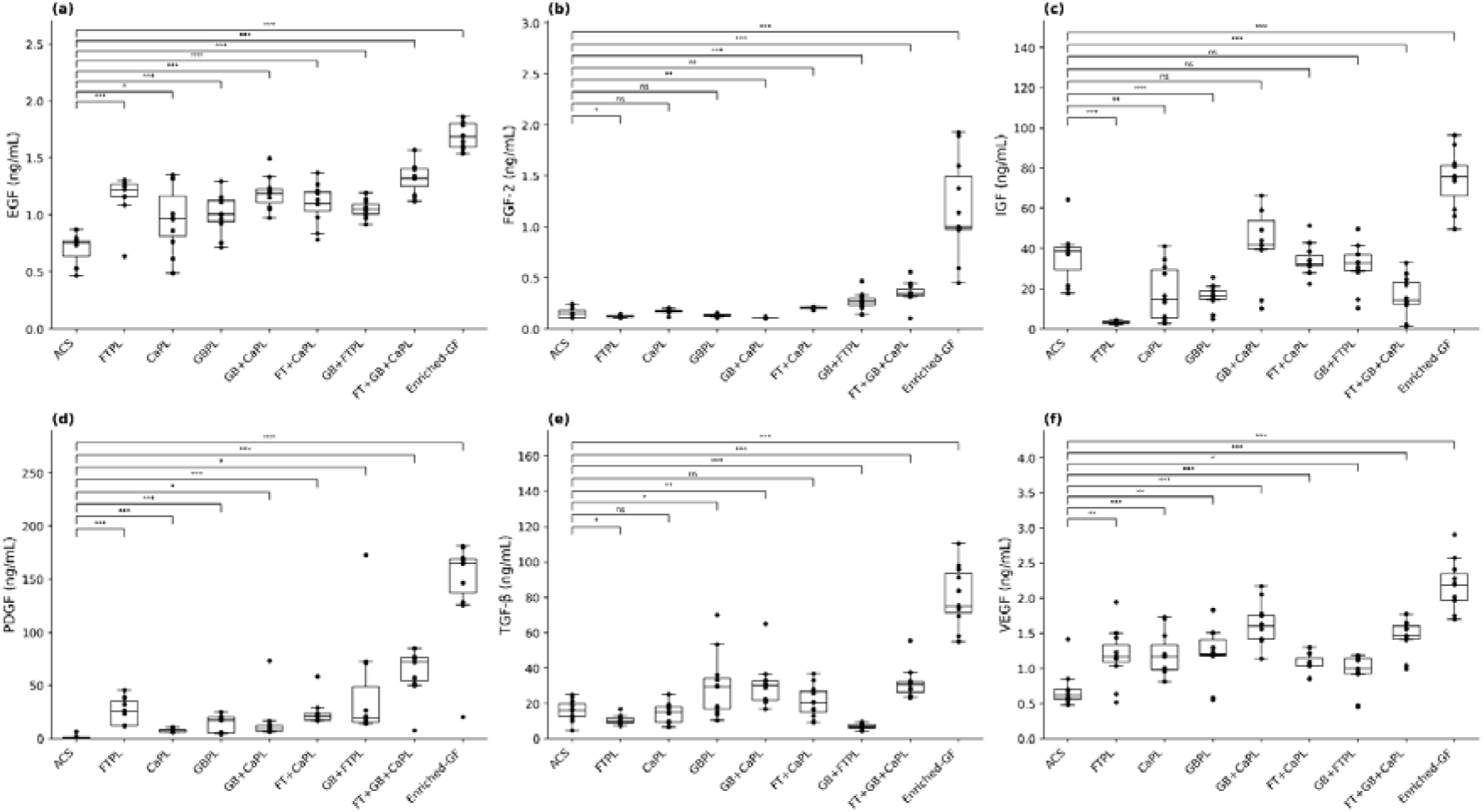
Soluble growth factor concentrations in different preparations. **A.** Epidermal Growth Factor (EGF), **B.** Fibroblast Growth Factor (FGF), **C.** Insulin-Like Growth Factor (IGF), **D.** Platelet-Derived Growth Factor (PDGF), **E.**Transforming Growth Factor Beta (TGF-B), and **F.** Vascular Endothelial Growth Factor (VEGF) protein levels were measured in duplicate by enzyme-linked immunosorbent assay (ELISA) in the platelet lysates of ten donors. ACS: autologous conditioned serum, FT-PL: freeze-thawed platelet lysate, Ca-PL: calcium-activated platelet lysate, GB-PL: glass bead platelet lysate, GB+CaPL: glass beads and calcium-activated platelet lysate, FT+Ca-PL: freeze-thawed and calcium-activated platelet lysate, GB+FTPL: glass beads and Freeze-thawed platelet lysate, FT+GB+Ca-PL: glass beads+freeze-thawed and calcium-activated platelet lysate and Enriched GF methods.Statistical significance was assessed using an independent samples t-test (*: P < 0.05, **: P < 0.01, ***: P < 0.001, ns: P ≥ 0.05).

**Table 1.**
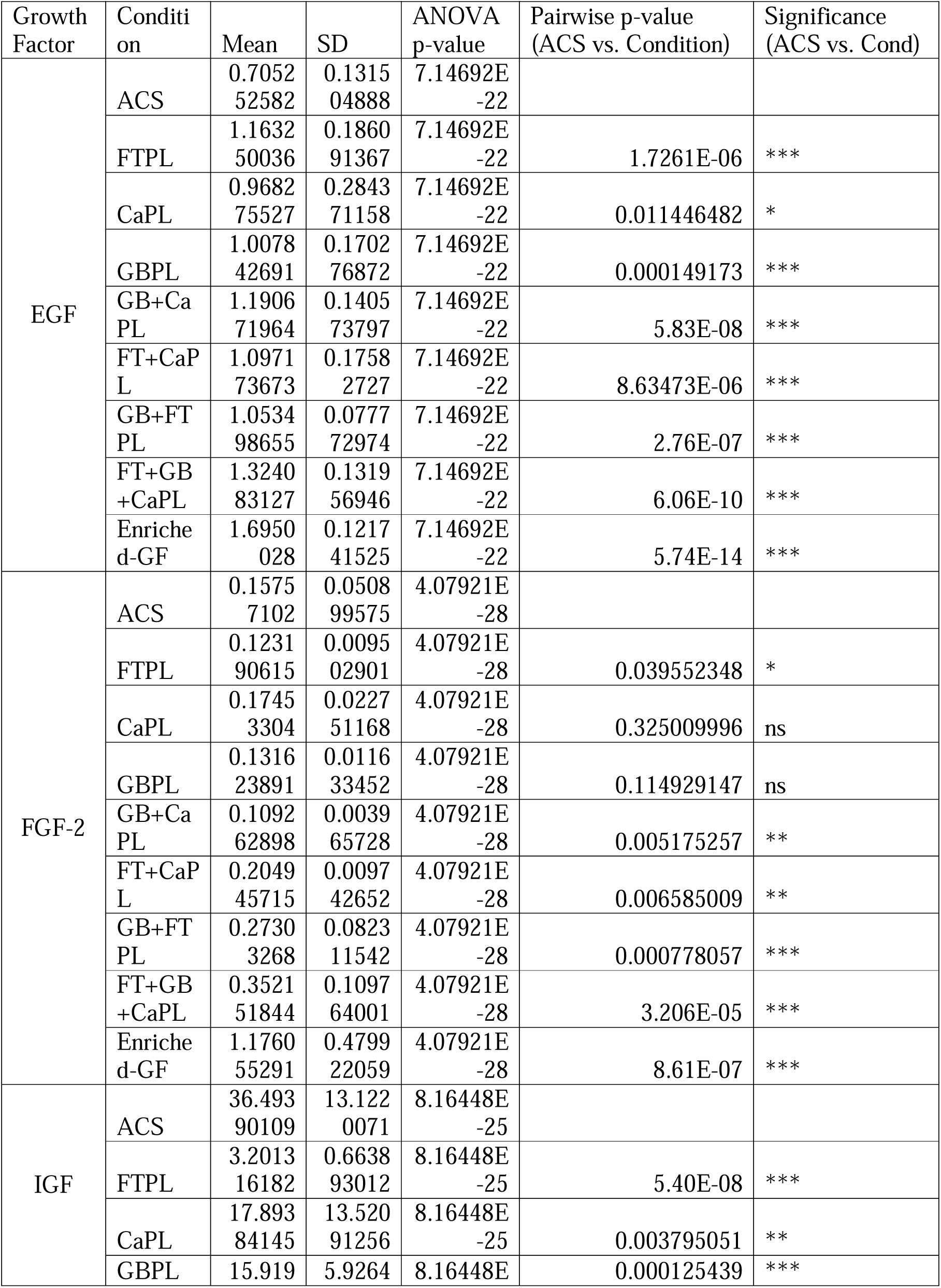
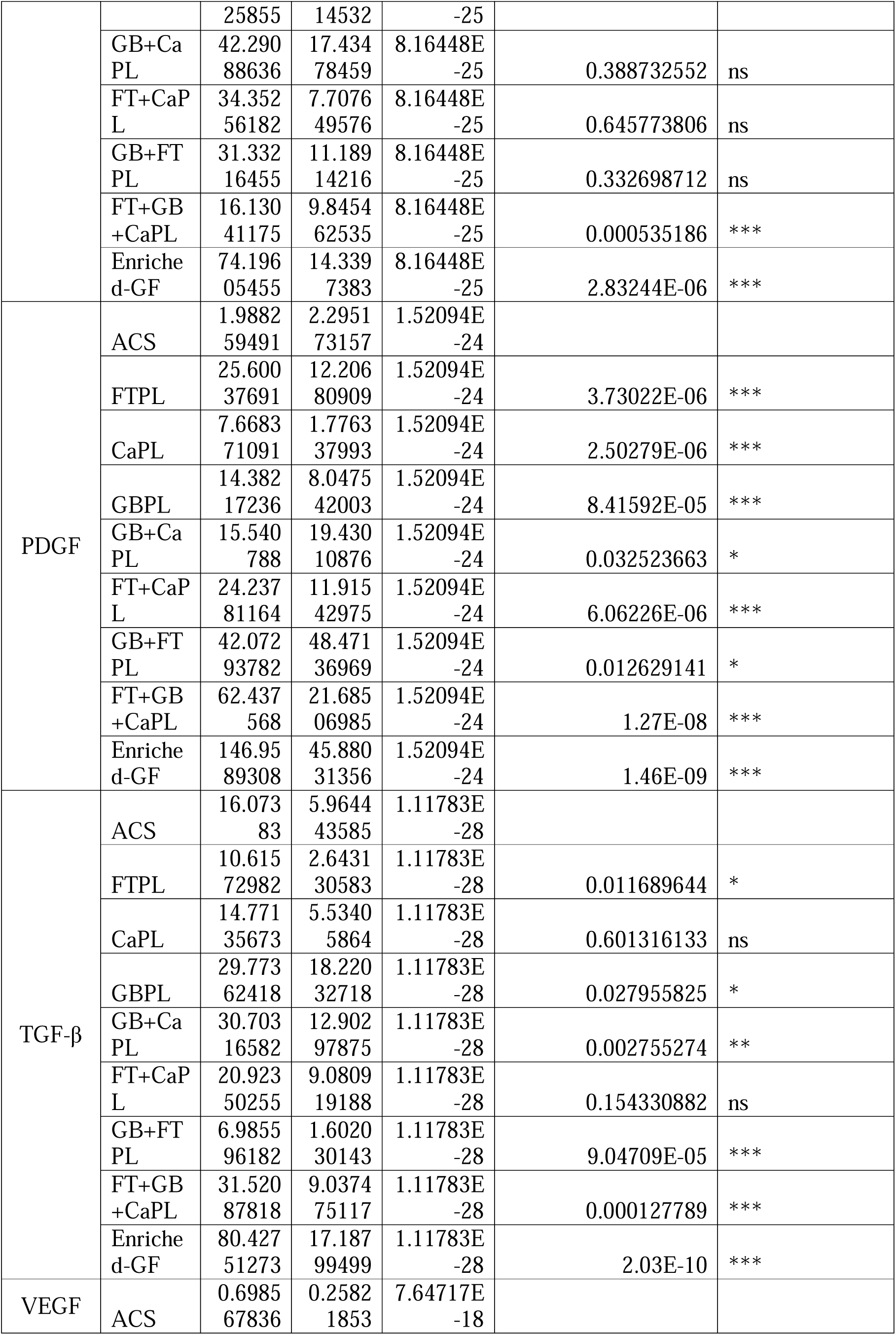
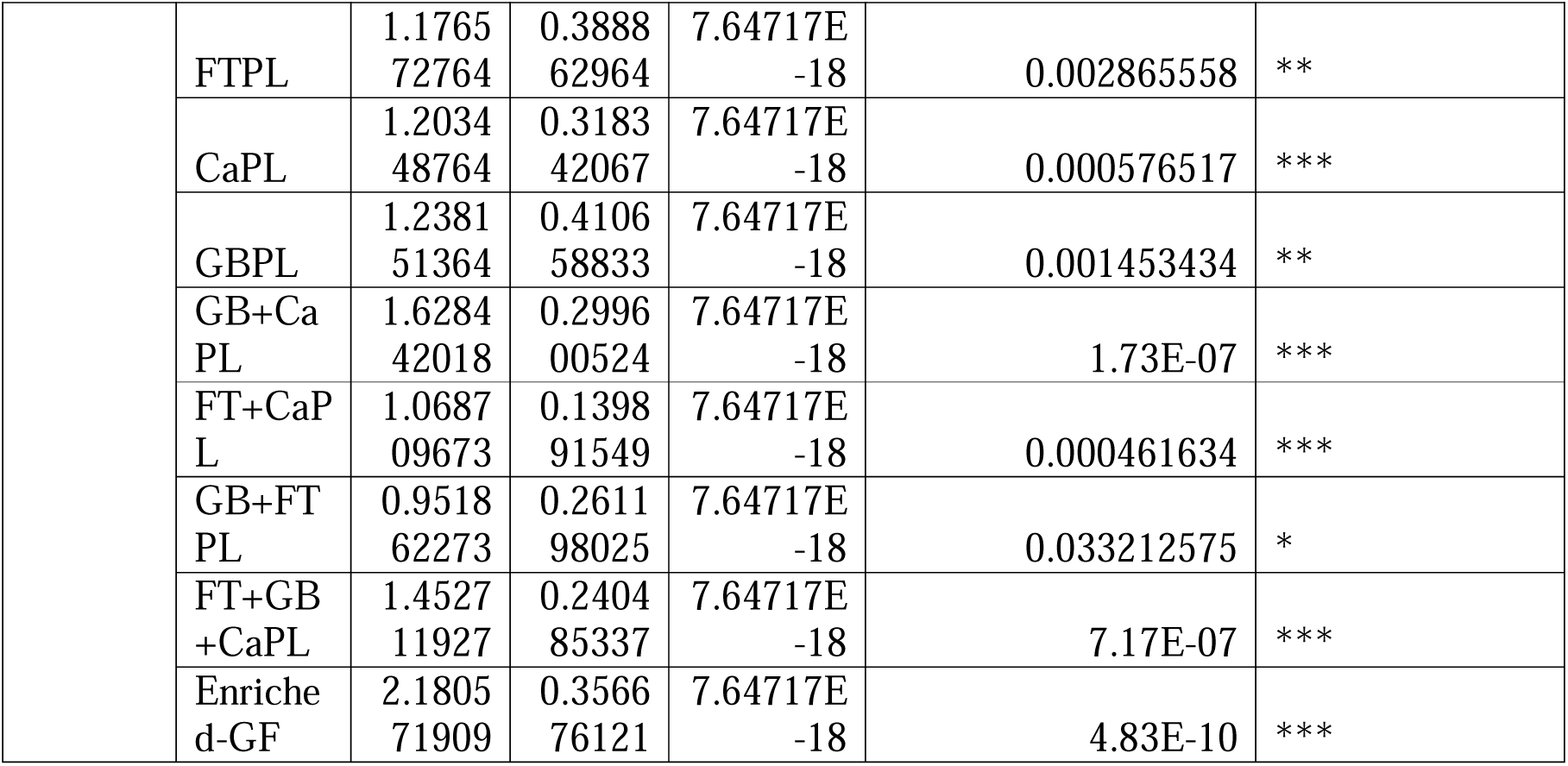

Collectively, these findings indicate that the multi-step activation of platelets—encompassing freeze–thaw disruption, Ca^2^-induced coagulation, and mechanical agitation of the fibrin matrix—results in a significantly enhanced and more consistent release of both regenerative and immunomodulatory growth factors. Consequently, the Enriched-GF protocol represents a standardized, high-yield platelet-derived biologic that effectively mitigates the variability limitations associated with traditional PRP and lysate preparation methods. Moreover, the reduced variability among donors suggests that the Enriched-GF protocol may offer greater consistency for both clinical and research applications. These results underscore the critical importance of optimizing activation methods to maximize the therapeutic efficacy of platelet-derived products.

## Discussion

The therapeutic potential of platelet-derived biologics is closely linked to the concentration and release kinetics of their bioactive GFs, both of which are determined by the platelet activation strategy employed. Among the newly developed blood-derived biologics, GFC and hPL have gained prominence as xeno-free, autologous alternatives to PRP. However, conventional methods for generating lysates often suffer from variability in platelet yield, incomplete degranulation, and protein degradation, frequently requiring heparin to prevent fibrin polymerisation—introducing additional biological and regulatory challenges ^24, 35^. The present study systematically compared several activation modalities and introduced a composite, additive-free approach that combines chemical, thermal, and mechanical activation. This multimodal strategy was designed to maximise GF release from platelets in a controlled ex vivo environment while improving yield stability and reproducibility.

The results demonstrate that Enriched-GF achieved superior growth factor release across all categories assessed, with PDGF and TGF-β exhibiting the highest absolute concentrations. PDGF levels increased more than fourfold relative to conventional single-activation approaches, highlighting the synergistic effect of combining FT–induced granule rupture, Ca-mediated thrombin activation, and GB–induced mechanical shear. This multimodal strategy recapitulates the sequential activation processes that occur during physiological platelet activation in vivo, enabling both rapid and sustained release of α-granule constituents.

Thermal-induced platelet lysis via freeze–thaw cycles initiates the immediate discharge of α-granule contents, producing the early PDGF burst ^26, 36^. In parallel, Ca-driven activation engages the thrombin–fibrin cascade, promoting fibrin crosslinking and supporting a prolonged secondary phase of PDGF release ^37^. Mechanical activation with glass beads provides contact-dependent stimulation, extending platelet degranulation and refining the temporal profile of PDGF secretion ^38, 39^. Consistent with these observations, Smith et al. reported that calcium activation generates early PDGF peaks, whereas ethanol prolongs release kinetics ^40^, confirming that multimodal activation strategies—such as Enriched-GF—couple rapid initial release (thermal/chemical) with controlled persistence (mechanical). This multi-phase activation closely mirrors in vivo platelet activation cascades and ensures both immediate and sustained growth factor availability, potentially accelerating wound repair and tissue regeneration

TGF-β release followed a similar trend, with the Enriched-GF method producing the highest concentrations across all conditions. Thermal activation accelerates the conversion of latent TGF-β complexes into their bioactive form by disrupting platelet membranes ^39^. Subsequent CaCl□-driven activation promotes fibrin matrix formation, which functions as a slow-release reservoir, preventing rapid diffusion and degradation. Mechanical stimulation via glass beads provides continued platelet engagement, extending degranulation and supporting sustained release over time. Together, these processes create a coordinated early surge and prolonged maintenance phase of TGF-β availability. This temporal profile is particularly relevant for wound healing, where TGF-β directs both the initial inflammatory response and later matrix remodelling. Consistent with this, Smith et al. noted that while some growth factors show a limited response to activation strategies ^40^, TGF-β is highly sensitive to combined physical–chemical cues due to its dependence on both extracellular matrix binding and regulated granule secretion.

IGF-1, a key mediator of metabolic activation, proliferation, and angiogenesis, also reached maximal levels under Enriched-GF. Thermal activation disrupts platelet membranes and partially cleaves IGF-binding proteins, thereby increasing the proportion of bioavailable IGF-1 ^41^. CaCl□-mediated chemical activation contributes to the initial peak of IGF-1 release, but this release may diminish without sustained stimulation. Mechanical activation appears to prolong platelet signalling, supporting sustained IGF-1 secretion over time. Consistent with this, cooling has been reported to inhibit IGF-1 release, underscoring the importance of controlled thermal cycling compared with low-temperature activation ^40^. Thus, the combined thermal rupture and chemical triggering characteristic of Enriched-GF may optimize IGF-1 kinetics in a manner well-suited for musculoskeletal tissue regeneration.

FGF-2, another factor crucial for cellular proliferation, metabolic support, and angiogenesis, similarly reached its highest concentrations under Enriched-GF. FGF-2 release is particularly sensitive to the mode of platelet disruption, as it exists both intracellularly and bound to heparan sulfate on platelet membranes ^42^. Thermal activation liberates cytoplasmic FGF-2, while CaCl□-driven fibrin formation serves to stabilize and protect the released protein from proteolysis. Mechanical shear from glass bead activation further sustains release over time, potentially prolonging its functional half-life. Prior work indicating that additives such as ethanol or mechanical agitation can modulate growth factor release kinetics without altering total yield aligns with this responsiveness of FGF-2 to activation modality ^43^. The dual thermal–chemical activation embedded in Enriched-GF therefore enhances both release and stability of FGF-2, supporting early angiogenic signaling and the subsequent proliferation of progenitor and stem cell populations.

EGF is critical in epithelial proliferation, tissue remodeling, and angiogenesis. Among all tested conditions, the highest EGF concentrations were observed with Enriched-GF. Thermal activation facilitates rapid release of EGF from α-granules, while CaCl□-mediated chemical activation supports simultaneous fibrin polymerization, helping to retain EGF in proximity to the platelet-derived matrix. Mechanical activation via glass beads provides ongoing platelet stimulation, sustaining a lower, prolonged release phase over time ^44^. Although activation modality does not uniformly enhance every growth factor, EGF appears particularly responsive to combined physical and chemical cues, with the most pronounced effects observed under multimodal activation ^40^. Thus, the triple mechanism within Enriched-GF likely enhances epithelial regenerative capacity by synchronizing an early release burst with a controlled, sustained delivery phase.

VEGF, a key driver of endothelial proliferation and neovascularization, also reached its highest concentrations under Enriched-GF, with comparably elevated levels observed in GB+Ca-PL samples. This pattern indicates a synergistic interaction between mechanical and CaCl□-mediated activation pathways. The consistent performance of mechanically engaged (GB-based) protocols suggests that mechanical shear supports more regulated VEGF release during platelet lysis, in contrast to conventional FTPL or WB + Ca protocols, which showed greater variability. Heat activation alone produces only modest VEGF release, while CaCl□-triggered thrombin activation can drive rapid but often transient degranulation ^40^. Mechanical activation introduces sustained membrane stimulation and surface contact, generating slower and more reproducible VEGF secretion ^37^. Although Smith et al. reported no major differences in VEGF yield across activation strategies, their data showing inhibitory effects of vitamin C underscore the advantage of purely physical–chemical modulation ^40^. The mechanical–chemical synergy embodied in Enriched-GF is therefore expected to stabilize platelet membranes and regulate VEGF kinetics, producing more uniform angiogenic signaling necessary for microvascular repair and tissue regeneration.

The superior performance of Enriched-GF, yielding growth factor concentrations exceeding reported ranges (PDGF: 80–160 ng/mL; TGF-β: 70–180 ng/mL; VEGF/EGF: 0.5–5 ng/mL), underscores the efficiency of synergistic activation compared with single-modality approaches. Previous studies by Delabie et al., Cavallo et al., and Smith et al. demonstrated partial gains using either calcium, freeze–thaw, or mechanical methods independently, yet the current results demonstrate that only their integration achieves complete platelet activation and maximised α-granule degranulation ^36, 40, 45^. The enhanced yields obtained here not only surpass the quantitative benchmarks of standard hPL preparations but also provide more consistent inter-donor reproducibility, reducing the biological heterogeneity that has hindered clinical translation.

Mechanistically, the stability observed across donor samples likely arises from CaCl□-mediated fibrin entrapment combined with GB-induced modulation of shear stress. This stabilising effect explains the reduced variability between donors and aligns with reports suggesting that combined physical–chemical activation provides more standardised GF output. The reproducibility of the Enriched-GF process, therefore, supports its suitability for integration into Good Manufacturing Practice (GMP)-compliant workflows, addressing one of the major limitations of existing platelet-derived products.

Comparative literature further contextualises these findings: while studies such as Oeller et al. and Cañas-Arboleda et al. highlighted the regenerative potential of hPL for mesenchymal stem cell expansion, their preparations suffered from inter-batch variability and dependency on additive stabilisers ^35, 46^. Similarly, Adhya et al. demonstrated the use of aged platelet concentrates for wound healing but noted limited growth factor recovery ^47^. By contrast, Enriched-GF produced higher and more stable concentrations of key GFs without the need for exogenous agents, providing a reproducible and donor-independent platform for research and clinical applications. The enriched angiogenic (VEGF, FGF-2) and mitogenic (PDGF, EGF) profiles observed here suggest particular suitability for musculoskeletal, dermal, and tissue regeneration.

Although the current study focused primarily on quantitative GF profiling, preliminary biological assays indicated enhanced proliferative potential of Enriched-GF preparations. Further work should assess their effects on angiogenesis, immunomodulation, and long-term tissue integration, alongside rigorous evaluation of storage stability and immunological safety. Nonetheless, these results establish a mechanistic and methodological framework for producing high-yield, reproducible, and additive-free platelet-derived products. The Enriched-GF protocol represents a next-generation strategy in regenerative biologics—combining simplicity, potency, and translational compatibility within a single, standardisable system.

## Conclusion

The therapeutic efficacy of platelet-derived products is predominantly attributed to paracrine signaling, wherein growth factors and cytokines released from platelet granules facilitate healing and regeneration. The limited extracellular survival of platelets highlights the necessity for technologies capable of efficiently isolating, stabilizing, and concentrating these soluble mediators in a standardized manner. Enriched-GF is a cell-free platelet-derived product that isolates growth factors from the patient’s own platelets without containing intact cells, thereby providing a standardized and reproducible preparation. In contrast to conventional human platelet lysate (hPL), Enriched-GF is xenofree, eliminating the need for Heparin, and being autologous, it poses no risk of immunogenicity or allergic reactions. When applied appropriately, Enriched-GF has the potential to expedite tissue repair, mitigate inflammation, and offer a safe and consistent therapeutic alternative for regenerative applications.

## Study Limitations

While the findings demonstrate clear advantages of the Enriched-GF protocol, several limitations must be acknowledged. First, the study was conducted on a relatively small donor cohort, which may limit statistical strength and generalizability; individual differences in platelet count, age, and plasma protein content could have influenced growth factor yield. Second, the study focused solely on biochemical quantification, without functional validation; thus, the biological activity of the released GFs in promoting cell proliferation, angiogenesis, or tissue repair remains to be confirmed. Third, GF stability and release kinetics beyond the immediate preparation window were not evaluated, and longer-term storage and shelf-life studies are required.

Future studies should include larger and more standardized donor groups, incorporate correlations between platelet concentration and release, extend release kinetics analysis, and perform in vitro and in vivo functional assays. Integrating clinically relevant tissue-repair models will be essential to determine whether the enhanced biochemical yields observed with the Enriched-GF method translate into superior therapeutic outcomes.

## Contributions

H.B. and M.S. designed and conducted the study. A.B. drafted the study protocol. A.B. and A.B. assisted in designing the protocol, supervised the study, and reviewed the results. H.B. contributed to writing, review and editing. J.L, M.F.V. and L.R. reviewed the results and the manuscript. H.B. and S.H.K. contributed to funding acquisition and project administration. All authors have read and agreed to the published version of the manuscript.

## Ethics approval and consent to participate Ethics Statements

This study was approved by the Institutional Committee for Stem Cell Research and Therapy (ICSCRT), Anupam Hospital, Rudrapur (Protocol No. AAH-03-2025, Version 2.0; approved 11 October 2024). Written informed consent was obtained from all participants.Written informed consent was obtained from all patients for participation in this study.

## Data Availability Statement

The data presented in this study are available on request from the corresponding author.

## Consent for Publication

Not applicable.

## Acknowledgements

We sincerely thank Lisa Fortier and Mary A. Ambach for their valuable suggestions and for reviewing the manuscript. We are also deeply grateful to the patients for their kind collaboration. The authors further acknowledge the laboratory staff for their assistance in the collection and processing of samples.

